# Patterns of host plant use do not explain mushroom body expansion in Heliconiini butterflies

**DOI:** 10.1101/2023.04.06.535866

**Authors:** Fletcher J. Young, Monica Monllor, W. Owen McMillan, Stephen H. Montgomery

## Abstract

The selective pressures leading to the elaboration of downstream, integrative processing centres, such as the mammalian neocortex or insect mushroom bodies, are often unclear. In *Heliconius* butterflies, the mushroom bodies are three to four times larger than their Heliconiini, and the largest known in Lepidoptera. Heliconiini lay almost exclusively on *Passiflora*, which exhibit a remarkable diversity of leaf shape, and it has been suggested that the mushroom body expansion of *Heliconius* may have been driven by the cognitive demands of recognising and learning the leaf shapes of local host plants. We test this hypothesis using two complementary methods: i) phylogenetic comparative analyses to test whether variation in mushroom body size is associated with the morphological diversity of host plants exploited across the Heliconiini; and ii) shape learning experiments using six Heliconiini species. We found that variation in the range of leaf morphologies used by Heliconiini was not associated with mushroom body volume. Similarly, we find interspecific differences in shape learning ability, but *Heliconius* are not overall better shape learners than other Heliconiini. Together these results suggest that the visual recognition and learning of host plants was not a main factor driving the diversity of mushroom body size in this tribe.

## Introduction

Patterns of investment in neural structures and pathways are shaped by the cognitive demands imposed by ecology, including foraging behaviours [1–3], mate detection and selection [4], and predator avoidance [5]. While increased investment in the sensory centres of the brain is largely driven by the sensory conditions experienced by an individual [6–10], the factors leading to the elaboration of downstream, integrative processing centres, such as the mammalian neocortex or insect mushroom bodies, are often less clear [11].

The Neotropical butterfly genus *Heliconius*, consisting of approximately 50 species, and nine closely related genera in the tribe Heliconiini (Lepidoptera: Nymphalidae), provides a novel and potentially powerful comparative framework for exploring the evolution of central brain structures [12]. In *Heliconius*, the mushroom bodies are markedly expanded, being three to four times larger than is typical for Lepidoptera, relative to brain size, including other Heliconiini genera ([13–15]). The mushroom bodies are historically studied as olfactory learning and memory centres in the predominant insect model species, *Drosophila melanogaster* [16–20]. However, across different taxonomic groups, mushroom bodies are innervated by a variety of sensory projection neurons, including visual and olfactory pathways, and the extent of innervation by different sensory modalities varies substantially across species [21]. Indeed, across insects, the mushroom bodies play a role in many visuals tasks including colour learning [22], visually-guided spatial navigation [23–25], reversal learning [26,27], and non-elemental associative learning [28].

Variation in the size and internal structure of the mushroom body likely reflects the relative importance of different sensory modalities in learnt behaviours and behavioural coordination, and selection for increased precision or fidelity of learnt associations. The expansion of the mushroom bodies in *Heliconius* therefore implies a major shift in information processing capacity in this genus, and the number of learnt patterns, or engrams, these butterflies can support [29]. The major expansion event in *Heliconius* co-occurs with the emergence of a dietary innovation unique amongst Lepidoptera – adult pollen feeding – and associated foraging specialisations (Figure 1) [30]. In collecting pollen, *Heliconius* establish ‘traplines’, foraging routes along which specific plants are regularly visited, suggesting a sophisticated capacity for spatial memory, probably using learnt visual landmarks [31–34]. The apparent cognitive demands of traplining have been hypothesised as driving the elaboration of the mushroom body in *Heliconius* [12–15]. Consistent with the explanation, the expansion of the mushroom bodies in *Heliconius* is primarily driven by increased visual, rather than olfactory input [15]. In addition, relative to the non-pollen-feeding *Dryas iulia, Heliconius erato* exhibit improved visual long-term memory and non-elemental learning, cognitive abilities which are important for the establishment and maintenance of traplines [15].

**Figure 1.**
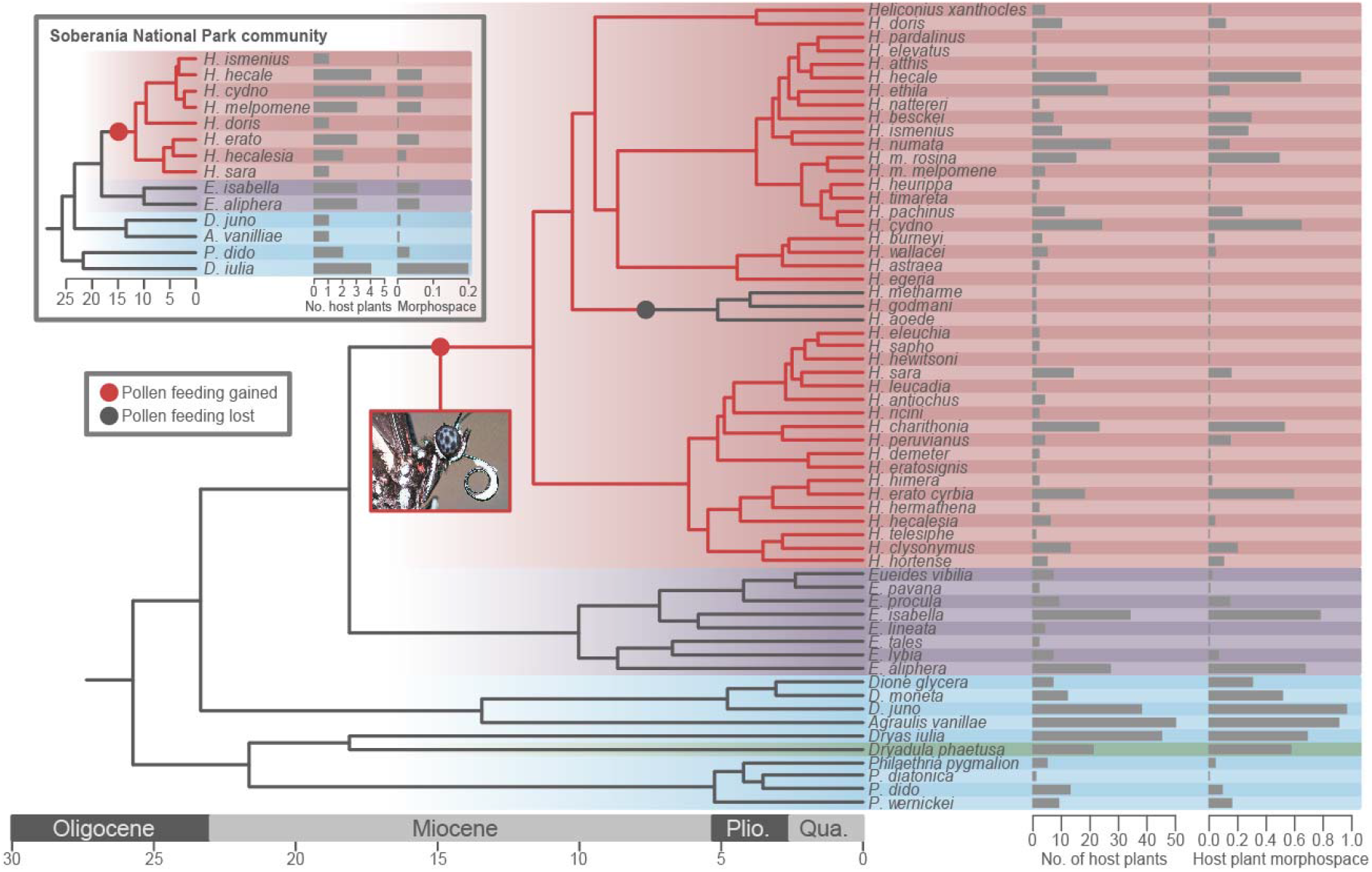
Host plant use across the Heliconiini showing number of host plants used, and the four-dimensional morphospace of their leaf shapes, quantified using Elliptical Fourier Descriptors. Background colours show phylogenetic shifts in the scaling relationship between the mushroom body and the rest of the central brain as identified in [15]. Inset shows host plant use in a specific Heliconiini community from Soberanía National Park, Panama. Tree adapted from [60].

However, although the largest shift in mushroom body size is observed in *Heliconius*, earlier bursts of mushroom body expansion are detected at the base of the *Heliconius-Eueides* clade, and independently in *Dryadula*. Additionally, the secondary loss of pollen feeding in *H. aoede* is not accompanied by a decrease in mushroom body size [15]. Thus, pollen feeding alone does not explain all variation in mushroom body size across the tribe. Increases in *Heliconius* mushroom body size have also been speculatively linked to the cognitive demands of identifying and learning host plants [35]. Herbivorous insects, which generally specialise on a limited range of plants, must solve the ecological challenge of identifying suitable host plants in complex, heterogenous environments [36]. Visually-oriented insects may identify host plants based on either a learned or innate “search image” for particular leaf shapes [37,38], with evidence for shape recognition in insects primarily deriving from studies on foraging bees [39–42]. However, within butterflies, *Eurema brigitta* exhibit a preference for leaves shaped like those of their larval host plant [43], while shape learning has been demonstrated in *Danaus plexippus, Battus philenor* and *Heliconius erato* in the context of oviposition and/or feeding [38,44–46].

*Heliconius* lay eggs almost exclusively on Passifloraceae, most commonly *Passiflora*, with varying degrees of specialisation. *Passiflora* display a remarkable diversity of leaf shape [47], and host plant use in *Heliconius* appears to be partly based on leaf shape recognition and learning through associative conditioning [46]. Leaf shape, therefore, seems to act as a long-distance cue for identifying suitable host plants, which are then confirmed by olfactory and gustatory signals at closer distances [46]. Given that many *Heliconius* species can lay on a range of *Passiflora*, there may have been selection for the ability to efficiently learn and remember leaf-shape search images for the suitable host plants species that are locally common. It has therefore been suggested that the diversity of *Passiflora* leaf shapes was driven by selection to escape visual identification of host plants by *Heliconius* [31,35,46,48]. Mushroom body expansion may, therefore, support a greater array of search images and enhanced shape-learning ability in *Heliconius*, facilitating improved visual identification of local host plants [35].

To date, shape learning has only been assessed in a single *Heliconius* species, *H. erato*, which was shown to learn shape information from both floral and host plant stimuli [46]. The degree to which this ability varies across species is therefore unclear. However, within butterflies, there is some evidence linking the mushroom bodies to host plant use and learning. When exposed to host plant environments of varying complexities, the relative generalist butterfly *Polygonia c-album* exhibited greater mushroom body plasticity than two relative specialists, *Aglais io* and *Aglais urticae* [49]. Similarly, in the cabbage white butterfly, *Pieris rapae*, larger mushroom body calyces were associated with an improved ability to locate difficult-to-learn red host plants, and experience with red hosts was positively related to increased mushroom body lobe size [50]. While previous analyses suggest mushroom body size in the Heliconiini is not explained by the number of host plants used by a species [15], it remains possible that generalist Heliconiini are targeting *Passiflora* species with morphologically similar leaves, meaning the number of host plant species used does not necessarily represent the diversity of search images used in host plant foraging.

Here, we therefore test whether mushroom body expansion in *Heliconius* is related to shape identification and learning using two complementary methods: i) phylogenetic comparative analyses to test whether variation in mushroom body size is positively associated with the morphological diversity of host plants exploited across the Heliconiini; and ii) comparative shape learning experiments across six Heliconiini species, using artificial feeders. We hypothesise that, if mushroom body expansion is linked to shape learning, *Heliconius* species would consistently outperform the *non-Heliconius* species in this task. Although our results suggest interspecific variation in the range of leaf morphologies used by Heliconiini, as well as interspecific variation in shape learning ability, we find no evidence that this explains the striking diversity of mushroom body size in this tribe.

## Methods

### i) Quantification of leaf morphospace and comparative analyses

We investigated whether the volumes of Heliconiini mushroom bodies vary with host plant use, controlling for the size of the rest of the central brain, using a recently published neuroanatomical dataset [15], combined with an established dataset on Heliconiini host plant use [51]. In total, 36 Heliconiini species had both host plant and neuroanatomical data. For these species, we used morphometric analysis to quantify the morphospace of host plant leaf shapes used by Heliconiini species, following [47]. Leaf images were sourced from the Global Plants database managed by JSTOR, the Encyclopedia of Life and the digital collections of the Muséum national d’histoire naturelle, Paris, and the Royal Botanic Gardens, Kew. These images were cropped to images of single leaves and edited to remove the petiole and non-leaf material (Figure S1). In total, we collected images for 686 leaves from 150 *Passiflora* species and 19 leaves from 11 *Dilkea* species.

We used *SHAPE* v 1.3 [52] to characterise leaf outlines with Elliptical Fourier Descriptors (EFDs), following established methods [47,53–57]. Using *SHAPE*, we binarized these images and performed a chain-code analysis (Figure S1) which we used to calculate normalised EFDs based on 40 harmonics [52]. We manually ensured that each leaf outline was consistently aligned. The R package *Momocs* v 1.3.2 was used to convert the resulting normalised EFD .nef file into a COE object, which was then subject to principal component analysis using *Momocs* [58]. With the R package *dispRity* v 1.6.0 [59] we used the first four principal components to calculate the four-dimensional volume of host plant morphospace exploited for each Heliconiini species. The four-dimensional morphospace volumes were then square-root transformed for all further analyses to better fit a normal distribution. We then conducted a series of phylogenetic GLMMs with *MCMCglmm*, using a recent phylogenetic tree [60] to test for relationships between host plant morphospace and the relative volumes of brain regions [61]. Controlling for allometric scaling with an independent measure of brain volume (rest of central brain, rCBR), we tested whether mushroom body volume varied the four-dimensional host plant morphospace, while also testing whether these relationships varied between *Heliconius* and *non-Heliconius* species. Since the mushroom body receives visual input, we also tested whether host plant morphospace explains volumetric variation in the medulla (the largest component of the visual pathway). Finally, because host plant use can vary between populations of a single Heliconiini species, it’s possible pooling data across populations skews the results by overestimating host plant use. We therefore also tested for these relationships within a single Heliconiini community (14 species) with well-described host plant use, from Gamboa, Panama, and the nearby Soberanía National Park [62]. All *MCMCglmm* models described above were run for 500,000 iterations, with a burn-in of 10,000 and a thinning factor of 500.

### ii) Shape-learning experimental protocol

To complement our comparative analyses, we performed shape learning assays in six Heliconiini species. All individuals used in the shape-learning experiments were reared from stocks established with locally caught, wild butterflies using the insectaries at the Smithsonian Tropical Research Institute (STRI) in Gamboa, Panama, in 2018. Stock butterflies were kept in 2×2×3 m mesh cages in ambient conditions with natural light. Larvae were reared in mesh pop-ups and provided with fresh leaves daily. *H. erato, Dryas iulia, Dryadula phaetusa* and *Agraulis vanillae* were reared on *P. biflora, H. melpomene* on *P. triloba*, and *H. hecale* on *P. vitifolia*. Training and testing of butterflies were conducted in 2×2×3 m mesh cages in ambient conditions under natural light. A single *Psychotria elata*, with all flowers removed, was placed in the rear right corner of these cages as a roosting site.

All individuals were freshly eclosed, to control for prior feeding experience. The day after eclosion, butterflies were transferred to a pre-training cage to familiarise them with the use of artificial feeders. Here, individuals were fed solely with red, circular feeders (Figure S2(a)), filled with a sugar-protein solution (20% sugar, 5% Vertark Critical Care Formula, 75% water, w/v) for one day. Artificial feeders were made from coloured foam with a centrally placed 0.5 ml Eppendorf tube. Throughout the experiment, all feeders were presented with a large, circular, green background so that the silhouette of the red shapes was clearly delineated in a consistent manner. After pre-training, butterflies were introduced to a testing cage to determine initial preference between two shapes – a “diamond” and a “star” shape (Figure S2(b)). The choice of these shapes was based on a previous study showing that *H. erato* can distinguish between them and tend to prefer star-shaped feeders over the diamond shape [46]. The testing cage contained 3 feeders of each shape, arranged randomly for each trial, and separated from each other by 15 cm (Figure S2(b)). To ensure that butterflies responded to visual cues only, feeders in the testing cages were empty. Preference testing lasted for four hours from 08:00 to 12:00 and was filmed using a GoPro Hero 5 camera mounted to a tripod. The film was then reviewed to count the number of feeding attempts per individual on each shape, with up to 40 attempts recorded. A feeding attempt was only counted if the butterfly landed on the feeder and probed it with its proboscis. Butterflies were then placed in a training cage for ten days which contained diamond-shaped feeders filled with sugar-protein solution and star-shaped feeders filled with an aversive saturated quinine solution. Through this combination of positive and negative stimuli we aimed to condition the butterflies to favour the diamond-shaped feeders over the star shapes. This training period lasted for 10 full days to provide ample opportunity for butterflies to learn [46], after which the trained feeding preferences were tested following the same protocol as the initial preference test.

### iii) Shape learning statistical analyses

Shape preferences and learning performance were analysed with generalised linear mixed models (GLMMs) using a binomial distribution with the *glmer* function from the package *lme4* v 1.1-27.1 in R v 4.1.2 [63]. Diagnostics for these GLMMs were assessed using the R package *DHARMa* v 0.4.4 [64]. All *post hoc* comparisons were made by obtaining the estimated marginal means using the R package *emmeans* v 1.7.0 and were corrected for selected multiple comparisons using the Tukey test [65]. Interspecific differences in initial shape preference were tested using a GLMM with species as a fixed effect and an individual-level random effect. For each species, we then tested whether initial preference towards a certain shape differed significantly from random using a GLMM with only individual-level random effects. Interspecific differences in shape learning performance were tested for using a GLMM with species and training as fixed effects, with an individual level-random effect. We also tested for an overall difference between *Heliconius* and non-*Heliconius* individuals using a GLMM with membership in *Heliconius* and training as fixed effects, with individual and species-level random effects. Sex was also initially included as a fixed effect in these models but was non-significant and so removed.

## Results

### i) Host plant generalism and the diversity of host plant morphologies

Principal component analysis of normalised EFDs based on 40 harmonics revealed that 93.3% of variation in *Passiflora* and *Dilkea* spp. host plant leaf shape is explained by the first four principal components, accounting for 60.8%, 17.8, 11.1% and 3.6% of variation, respectively (Figure 2). There was a significant positive relationship between the number of host plants recorded as being used by a species and the leaf-shape morphospace covered by those plants (F-value=199.8, d.f.=58, P<0.0001, R^2^ = 0.775, Figure 3(a)). However, over 20% of the variance in morphospace of host plants used by *Heliconius* was not explained by the number of plants used by the given *Heliconius* species. Several marked outliers further demonstrate the value of accounting for phenotypic diversity of host plants in characterising the search image landscape of a butterfly species (Figure S2). Overall, the morphospace volumes of host plants used by *Heliconius* did not significantly differ from other Heliconiini (Figure 1; pMCMC = 0.120). Comparing between Heliconiini genera, *Agraulis* host plants have a significantly larger morphospace than those of *Eueides, Heliconius* and *Philaethria* (Table S1), but these differences are vulnerble to correction for multiple comparisons (Table S2).

**Figure 2.**
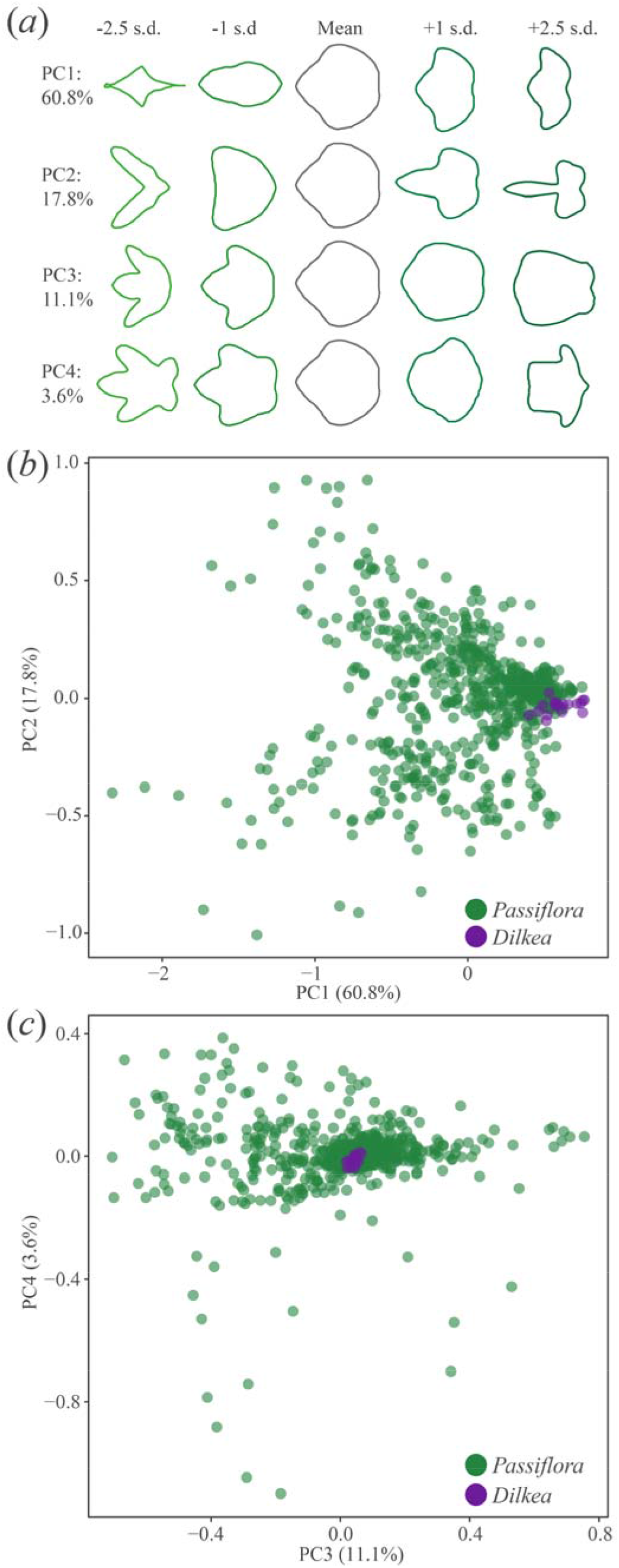
Principal components (PC) accounting for shape variation in Heliconiini host plants (150 *Passiflora* and 10 *Dilkea* species) characterised using Elliptical Fourier Descriptors. (*a*) ‘Eigenleaf representations at ±1 and ±2.5 SD for the first four PCs, with percentage of variance explained for each. (*b*) PC1 vs PC2. (*c*) PC3 vs PC4. Each point represents a single leaf.

**Figure 3.**
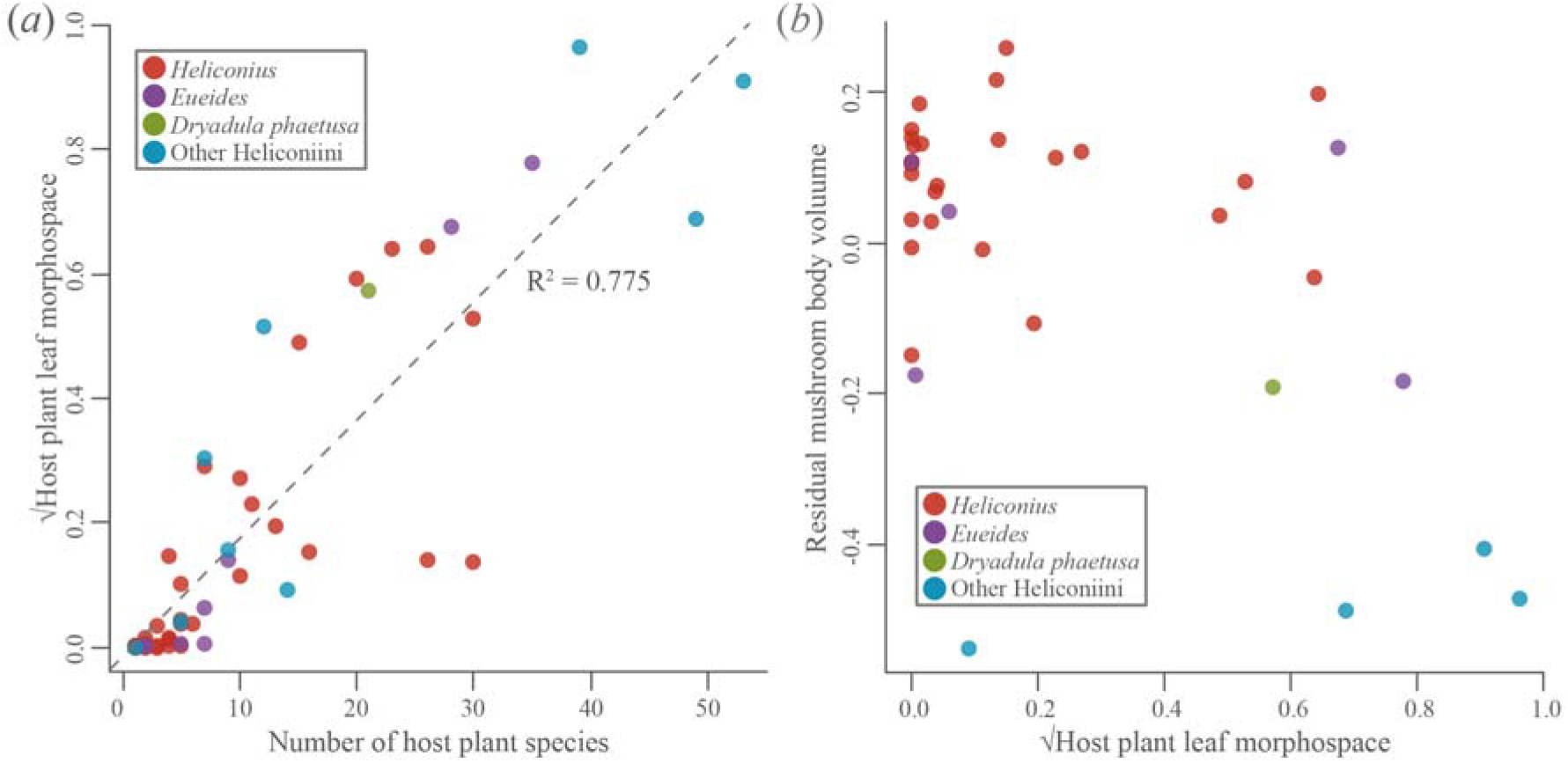
*(a)* Across the Heliconiini, host plant number is significantly positively correlated with the leaf shape morphospace of those plants. Dashed line shows linear regression (F-value=199.8, d.f.=58, P<0.0001, R^2^=0.775). *(b)* Relative mushroom body size (controlling for the size of the rest of the central brain) is not significantly associated with host plant leaf morphospace for either *Heliconius* or the outgroup Heliconiini.

### ii) Diversity of host plant use not associated with mushroom body size in Heliconiini

Across all Heliconiini, we found no relationship between mushroom body size and host plant morphospace (pMCMC = 0.214; Figure 3(b)). This was true both within *Heliconius* (pMCMC = 0.876), and among *non-Heliconius* Heliconiini (pMCMC = 0.391) separately. We further explored whether investment in the medulla, the largest neuropil of the optic lobe, could be associated with host plant use. Again, we did not detect an association between medulla volume and host plant morphospace in all Heliconiini (pMCMC = 0.818), *Heliconius* (pMCMC = 0.964), or the outgroup Heliconiini (pMCMC = 0.940) individually.

Finally, we repeated our analyses within a specific Heliconiini community of 14 species (Figure 1) around Gamboa, Panama, and the Soberanía National Park [62]. Nine *Passiflora* species are known to be used by this community, with butterflies ranging from specialists laying on a single species to relative generalists laying on five species. Neither host plant number (pMCMC=0.313) nor leaf shape diversity (pMCMC=0.309) was significantly associated with investment in mushroom body size.

### ii) No evidence Heliconius are better shape learners than other Heliconiini

In our shape learning experiment, we assayed 162 individuals from 6 species with well characterised host plant use. All six species showed a significant initial preference for the star-shape feeders over the diamond feeders, with stars accounting for approximately two-thirds of all feeding attempts (Figure 4, Table S3). Initial shape preference did not differ between species (χ^2^=2.709, d.f.= 5, *P*=0.746). Learning ability, however, varied significantly between species (χ^2^=18.565, d.f.= 5, *P*=0.002; Figure 4), but *Heliconius* as a whole were not superior shape learners than the other Heliconiini (χ^2^=0.128, d.f.= 1, *P*=0.720; Figure 4). Rather, shape learning ability was scattered across the phylogeny. *Agraulis vanillae, Dryadula phaetusa, H. melpomene* and *H. erato* showed a significant shift in preference towards the diamond shape after training, while *Dryas iulia*, and *H. hecale* showed no shift (Table S4, Figure 4). Notably, even the species that did show a significant shape-learning effect only showed a slight shift in preference and still maintained a bias towards stars over diamonds (Figure 4). This low effect size, which is consistent with previous experiments in *H. erato* [46], suggests that increasing the difficulty of the shape-learning task would not reveal a difference in shape-learning ability between *Heliconius* and the outgroup Heliconiini. We also note that visual acuity is unlikely to explain these results as naïve butterflies from all species show a similar preference for the star-shape, suggesting that all species are capable of distinguishing between these shapes.

**Figure 4.**
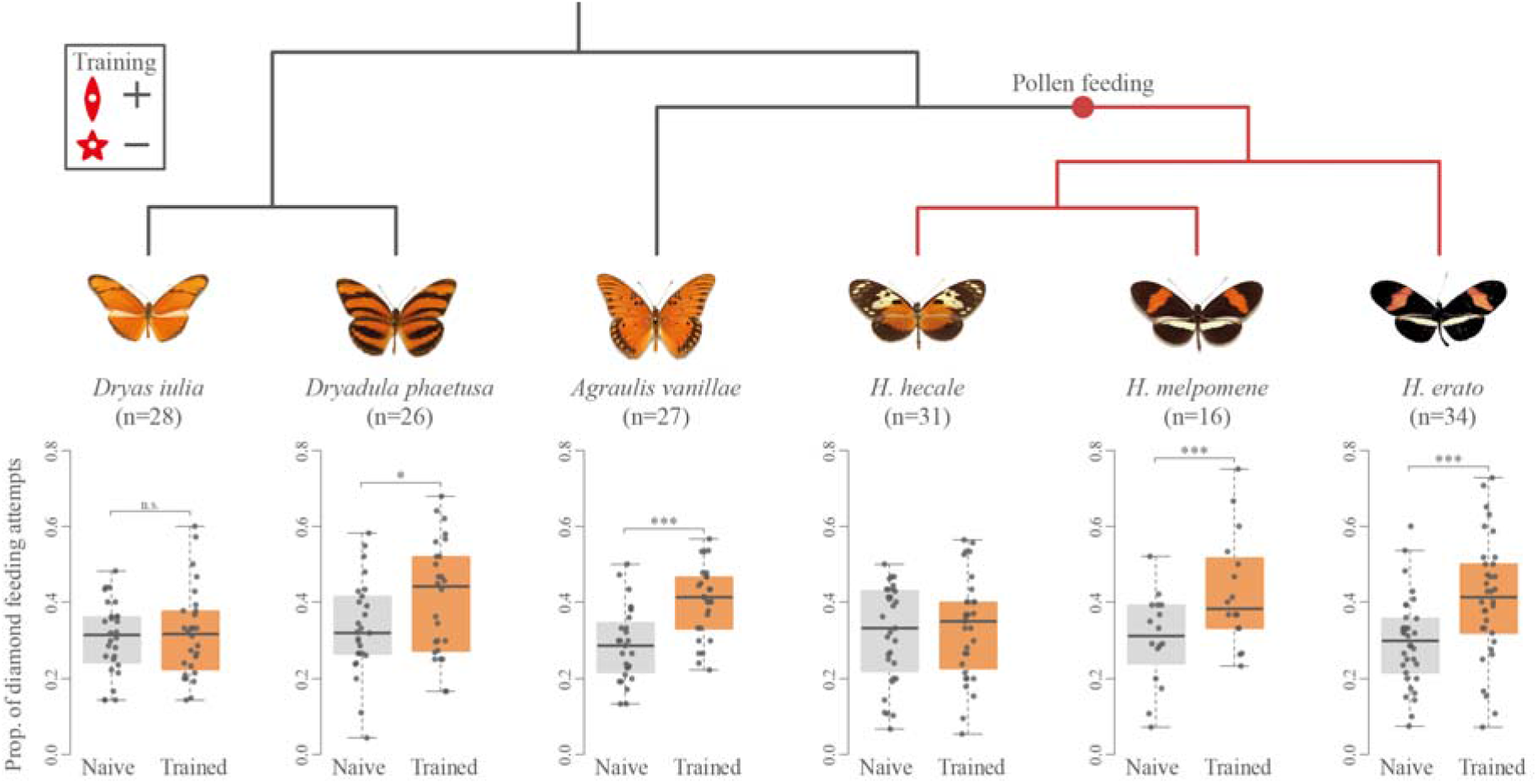
Shape learning in six Heliconiini species. Boxplots show frequency of feeding on diamond shapes when given a choice between diamond and star shapes, before and after 10 days’ training. Shape learning performance varies significantly between species, but *Heliconius* are not better shape learners overall. * = P<0.05, *** = P < 0.001.

## Discussion

*Heliconius* butterflies exhibit a dramatic expansion of the mushroom bodies, which has been hypothesised to have been driven by the cognitive demands of trapline foraging for pollen [15]. However, separate instances of mushroom body expansion in non-pollen-feeding Heliconiini suggest that pollen feeding does not fully explain this variation. Another hypothesis posits that mushroom body expansion in *Heliconius* is related to the cognitive demands of identifying suitable host plants [35], which relies, in part, on leaf shape learning and recognition [46]. We tested this hypothesis using a combination of phylogenetic comparative methods to analyse neuroanatomical and host plant morphospace data. Further, we tested shape learning ability in six Heliconiini species.

It was previously found that, within the Heliconiini, mushroom body size is not associated with host plant number [15]. However, our results show that the diversity of host plant leaf shapes is largely, but not fully, explained by host plant number (Figure 3(a)). Our first test of the host plant hypothesis, therefore, used phylogenetic comparative analyses to test whether host plant morphospace explained any variation in mushroom body size across the Heliconiini. Our results showed no evidence of an association between mushroom body size and the leaf shape diversity of a species’ host plants (Figure 3(b)). Additionally, interspecific variation in medulla volume, the largest visual neuropil, was also not associated with host plant morphospace. This suggests that the visual learning and recognition of host plants was likely not an important driver of the several incidences of mushroom body expansion seen in the Heliconiini [15]. This is somewhat surprising given the apparent role of visual cues in *Heliconius* host plant selection [46] and the previous data implicating the mushroom bodies in the host plant foraging in the butterfly *Pieris rapae* [49,50].

However, host plant use can vary between populations for some Heliconiini species [66], and *Passiflora* diversity within a given area can be limited to approximately 10 species [31]. Therefore, for some Heliconiini populations, host plant number, and their estimated leaf shape diversity, is likely overestimated for some species. We further accounted for this potential discrepancy by analysing host plant use within a single, well studied community of Heliconiini near Gamboa, Panama, within the Soberanía National Park [62]. The results of this analysis reflected the results of the wider analysis, also showing no relationship between differences in mushroom body size and host plant number or leaf shape diversity. These results suggest that the identification of host plants based on leaf shape has not been an important factor influencing brain evolution in the Heliconiini, at least at a volumetric scale. Indeed, while shape cues are used by butterflies to detect host plants [38,43,44,46], olfactory cues are also important, although these are probably used at shorter distances [36]. Indeed, in a previous study, although *H. erato* trained on a certain host plant leaf shape approached that shape more frequently, the total landings and number of eggs laid did not differ between shapes [46].

As a second test of the host plant hypothesis, we tested shape learning ability in three *Heliconius* and three *non-Heliconius* Heliconiini species from the Panamanian community. The six species assessed in this study exhibited a clear ability to distinguish between shapes, all showing a marked preference for star-shaped feeders over diamonds, consistent with previous experiments showing a similar bias in *H. erato* (Figure 4) [46]. This preference is likely explained by the resemblance of the star-shape to the radial symmetry of the flowers typically exploited by these Heliconiini [31,67,68]. Visual cues, including “search images”, therefore appear to play an important role for Heliconiini in identifying key resources in their environment. Notably, naïve Heliconiini possess an inherent attraction towards certain shapes, similar to previous findings in honeybees [42] and the butterflies *Battus philenor* and *Eurema brigitta* [38,43].

Our results also corroborate previous findings of shape learning in *H. erato* [46], while also demonstrating comparable abilities in *H. melpomene* and the non-*Heliconius* Heliconiini *Agraulis vanillae* and *Dryadula phaetusa* (Figure 4). However, we also found significant variation in shape-learning performance between species, with neither *H. hecale* and or *Dryas iulia* showing evidence of having learned the shape cue. Contrary to our prediction however, *Heliconius*, as a group, were not superior shape learners than the *non-Heliconius* species (Figure 4). The lack of difference in shape-learning performance between *Heliconius* and *non-Heliconius* Heliconiini, suggests that mushroom body expansion in *Heliconius* is not associated with enhanced shape learning ability. This result casts doubt over the hypothesis that the expanded mushroom bodies of *Heliconius* are adapted for shape learning related to the visual recognition of *Passiflora* host plants.

In addition, even amongst the Heliconiini species which did exhibit a significant shape learning effect, the frequency of feeding attempts on diamond shapes remained below 50% (Figure 4). The innate preference for the star shape appears difficult to alter through associative conditioning, even when the star shape is presented with an aversive stimulus. This suggests that visual identification of resources in Heliconiini butterflies may be driven more by innate search images than learned shapes. One important caveat, however, is that to test shape learning in an interspecific, comparative framework, we conducted the experiment in a feeding, rather than oviposition context. The Heliconiini involved in this study use of a range of different host plants with varying leaf shapes and using the same shape cues for each butterfly species would not have been possible in an oviposition context. Additionally, testing the learning abilities of butterflies in a feeding context also has the advantage of far greater tractability, permitting the sample sizes necessary for a comparative behavioural experiment across six species. In some cases the importance of different types of cues can be context dependent [69–71]. However, the shape learning abilities of *H. erato* translate across both feeding and oviposition contexts [46], and a similarly fixed preference to the shape preference we observe was seen in the selection of host plants by *H. erato phyllis*, which was not able to be altered by experimental conditioning [66]. This contrasts with the apparent ease with which Heliconiini can learn, and modify, colour associations [72], a contrast also observed in monarch butterflies [73] and honeybees [74,75] (but not, interestingly, in the Hymenopteran parasitoid *Microplitis croceipes* [76]). We therefore suggest that shape-learning performance in a feeding context can be interpreted as a measure of general shape-learning ability in Heliconiini butterflies.

Combining the shape-learning experiment with comparative analyses of host plant diversity provides a complementary, alternative test of our overriding hypothesis. Together, our results do not support the hypothesis that visual identification of host plants based on leaf shape has been an important factor in driving MB expansion in *Heliconius*. The cognitive demands of traplining, therefore, remains as the most likely selective pressure, driving mushroom body expansion in *Heliconius* butterflies. Nevertheless, it is possible that *Passiflora* plants are incorporated into *Heliconius* pollen traplines [31], although this has not been formally observed. Further data, both from behavioural experiments, including assessments of spatial learning ability, and more comprehensive field observations of traplining behaviour across a wide sampling of Heliconiini species are necessary to directly test these hypotheses.

