## Supplementary materials for "Patterns of host plant use do not explain mushroom body expansion in Heliconiini butterflies"

**Supplementary material**

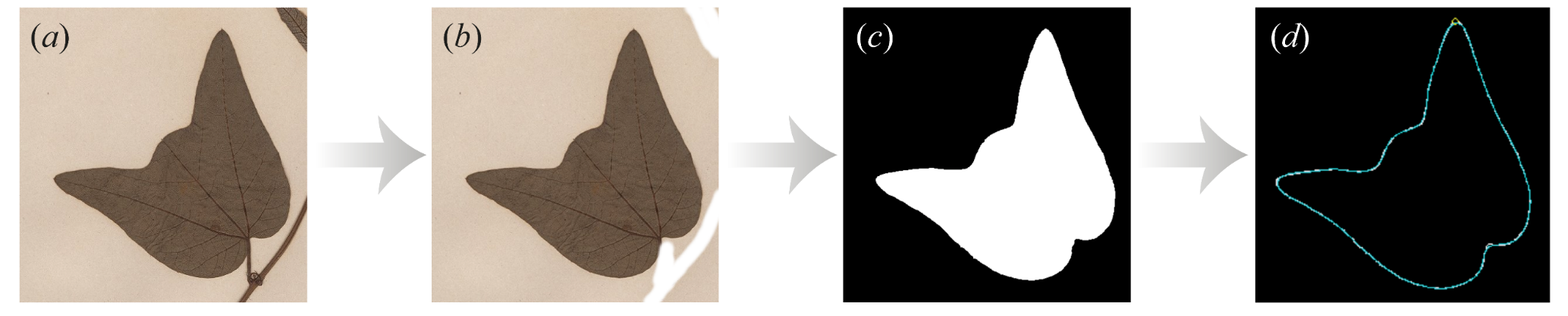

**Figure S1.** Image processing pipeline for chain code analysis in SHAPE, showing a Passiflora capsularis leaf. (a) Cropped, raw image of single leaf. (b) Manual removal of petiole and extra material. (c) Image is binarized. (d) Chain code analysis of the shape outline.

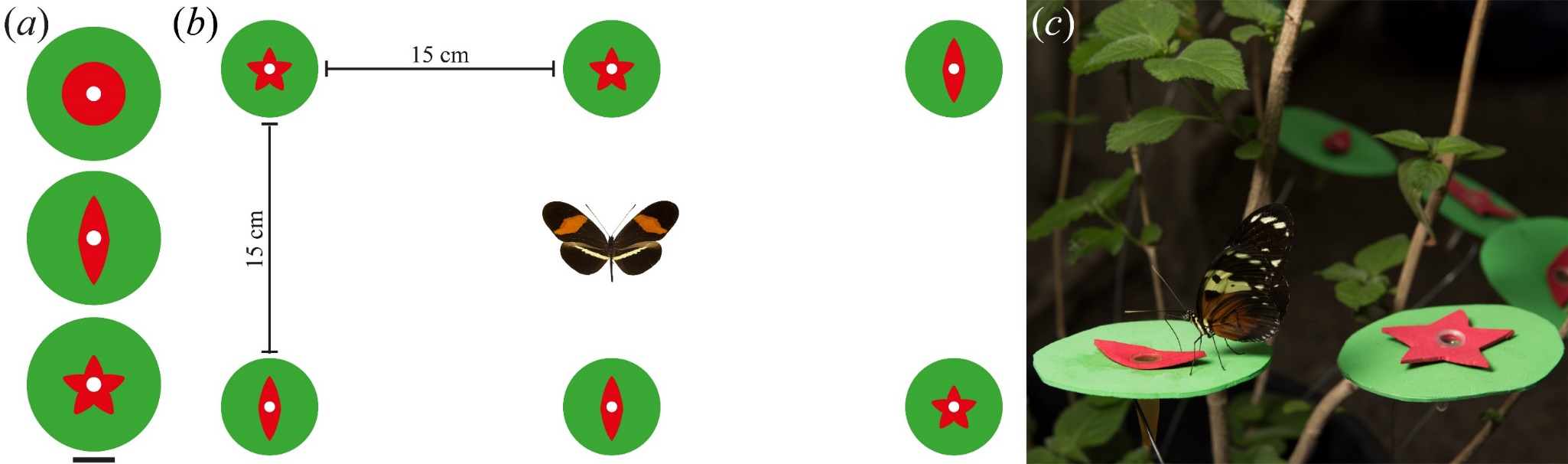

**Figure S2.** (a) The types of feeders used in the shape learning experiment. Circular feeder was used in pre-training to accustom butterflies to using the feeders. Butterflies were training to associate a food reward with the diamond-shaped feeders and a aversive quinine solution with the star shape. Scale bar = 20 mm. (b) Example of feeder arrangement during preference trials. Feeders were evenly spaced but randomly arranged for each trial. (c) Training environment with Heliconius hecale and the two feeder shapes.

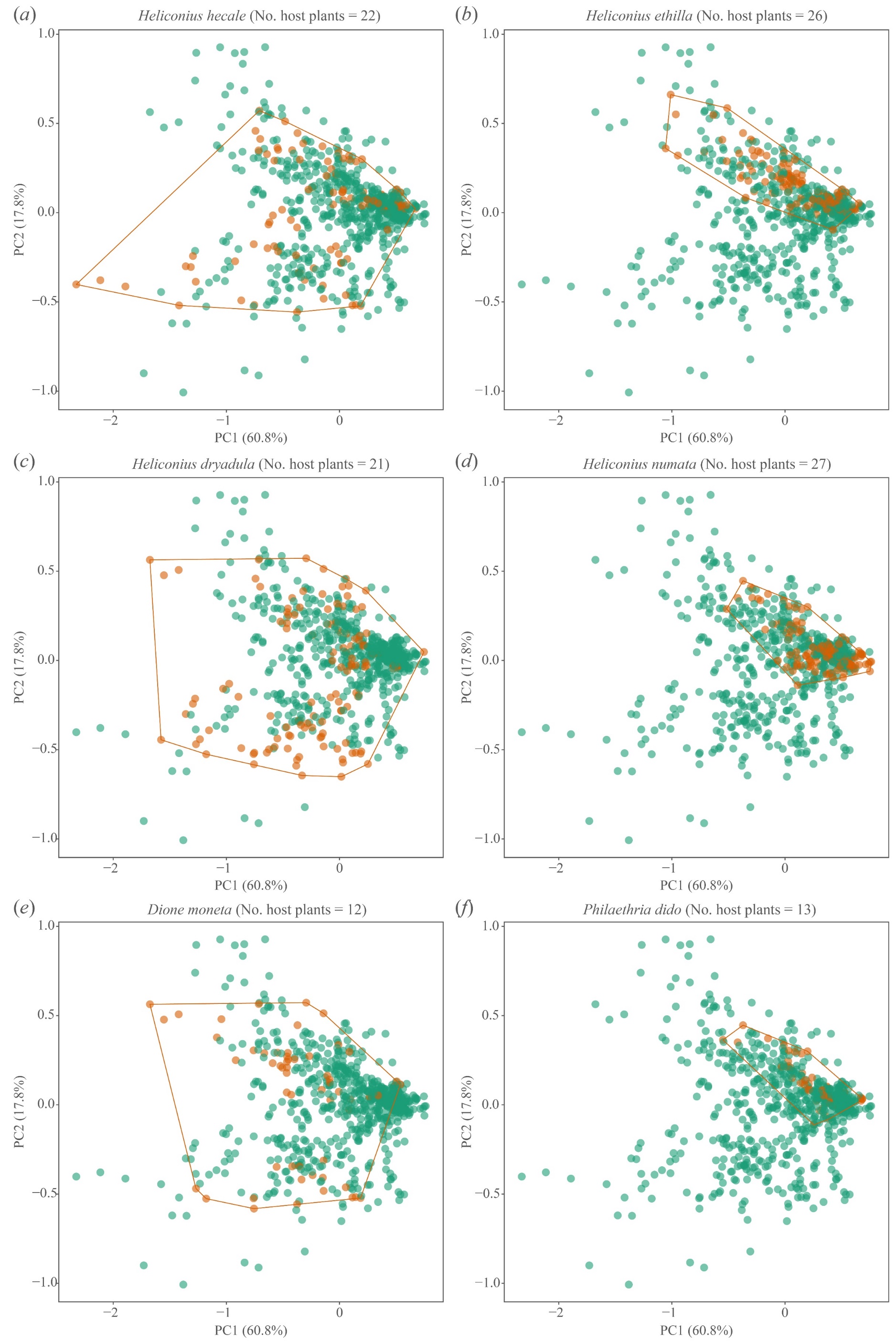

**Figure S3.** 2-dimensional host plant leaf morphospace for six Heliconiini species illustrating the difference between host plant number and disparity of host plant leaf shape. Species on the left use less host plant species but cover a greater leaf shape morphospace than those on the right. Leaves used by a given species shown in orange. Each point represents a single leaf sample.

**Table S1.** Uncorrected pairwise comparisons between Heliconiini genera in the morphospace volume of host plants used, characterised using Elliptical Fourier Descriptors. P-values shown above the diagonal and estimated pairwise differences below. * = P<0.05.

|  | ***Philaethria*** | ***Dryadula*** | ***Dryas*** | ***Agraulis*** | ***Dione*** | ***Eueides*** | ***Heliconius*** |
| --- | --- | --- | --- | --- | --- | --- | --- |
| ***Philaethria*** |  | 0.066 | 0.063 | 0.013* | 0.066 | 0.596 | 0.912 |
| ***Dryadula*** | 0.502 |  | 0.784 | 0.411 | 0.932 | 0.307 | 0.165 |
| ***Dryas*** | 0.605 | -0.103 |  | 0.551 | 0.838 | 0.163 | 0.071 |
| ***Agraulis*** | 0.841 | 0.339 | 0.236 |  | 0.324 | 0.036* | 0.012* |
| ***Dione*** | 0.533 | 0.031 | -0.072 | 0.308 |  | 0.162 | 0.052 |
| ***Eueides*** | 0.146 | 0.356 | 0.459 | 0.695 | 0.387 |  | 0.537 |
| ***Heliconius*** | 0.028 | 0.474 | 0.577 | 0.813 | 0.506 | 0.118 |  |

**Table S2.** Pairwise comparisons between Heliconiini genera in the morphospace volume of host plants used, characterised using Elliptical Fourier Descriptors and corrected for multiple comparisons using Tukey’s test. P-values shown above the diagonal and estimated pairwise differences below.

|  | ***Philaethria*** | ***Dryadula*** | ***Dryas*** | ***Agraulis*** | ***Dione*** | ***Eueides*** | ***Heliconius*** |
| --- | --- | --- | --- | --- | --- | --- | --- |
| ***Philaethria*** |  | 0.735 | 0.509 | 0.169 | 0.520 | 0.998 | 1.000 |
| ***Dryadula*** | 0.502 |  | 1.000 | 0.983 | 1.000 | 0.949 | 0.808 |
| ***Dryas*** | 0.605 | -0.103 |  | 0.997 | 1.000 | 0.804 | 0.543 |
| ***Agraulis*** | 0.841 | 0.339 | 0.236 |  | 0.957 | 0.357 | 0.149 |
| ***Dione*** | 0.533 | 0.031 | -0.072 | 0.308 |  | 0.802 | 0.452 |
| ***Eueides*** | 0.146 | 0.356 | 0.459 | 0.695 | 0.387 |  | 0.996 |
| ***Heliconius*** | 0.028 | 0.474 | 0.577 | 0.813 | 0.506 | 0.118 |  |

**Table S3.** Naïve shape preference for each species, tested using a generalised linear mixed model with individual-level random effect. All species significantly preferred star-shaped feeders over diamonds. *** = P<0.001.

| **Species** | **z value** | **P-value** | **Preferred shape** |
| --- | --- | --- | --- |
| *Dryas iulia* | -9.368 | <0.0001*** | Star |
| *Dryadula phaetusa* | -6.497 | <0.0001*** | Star |
| *Agraulis vanillae* | -10.04 | <0.0001*** | Star |
| *H. hecale* | -7.045 | <0.0001*** | Star |
| *H. melpomene* | -5.912 | <0.0001*** | Star |
| *H. erato* | -8.912 | <0.0001*** | Star |

**Table S4.** Pairwise comparisons between untrained and trained shape preferences for each species based on the generalised linear mixed model with training and species as fixed effects and an individual-level random effect, corrected for multiple comparisons using Tukey’s test. * = P<0.05; ** = <0.01; *** = P<0.001.

| **Species** | **z ratio** | **P-value** |
| --- | --- | --- |
| *Dryas iulia* | -0.629 | 0.989 |
| *Dryadula phaetusa* | -2.883 | 0.023* |
| *Agraulis vanilla* | -4.717 | <0.0001*** |
| *H. hecale* | -0.869 | 0.946 |
| *H. melpomene* | -3.822 | 0.0008*** |
| *H. erato* | -4.949 | <0.0001*** |
